# Global coordination of protrusive forces in migrating immune cells

**DOI:** 10.1101/2024.07.26.605242

**Authors:** Patricia Reis-Rodrigues, Nikola Canigova, Mario J. Avellaneda, Florian Gaertner, Kari Vaahtomeri, Michael Riedl, Jack Merrin, Robert Hauschild, Yoshinori Fukui, Alba Juanes Garcia, Michael Sixt

## Abstract

Efficient immune responses rely on the capacity of leukocytes to traverse diverse and complex tissues. To meet such changing environmental conditions, leukocytes usually adopt an amoeboid configuration, utilizing their forward-positioned nucleus as a probe to identify and follow the path of least resistance among pre-existing pores. We show that in dense environments, where even the largest pores preclude free passage, leukocytes switch polarity and position their nucleus behind centrosome and organelles. In this mesenchymal configuration, local compression of the cell body triggers assembly of a central F-actin pool, located between cell front and nucleus. Central actin pushes outward to transiently dilate a path for organelles and nucleus. Pools of central and front actin are tightly coupled and experimental depletion of the central pool enhances actin accumulation and protrusion formation at the cell front. Although this shifted balance speeds up cells in permissive environments, migration in restrictive environments is impaired, as the unleashed leading edge dissociates from the trapped cell body. Our findings establish an actin regulatory loop that balances path dilation with advancement of the leading edge to maintain cellular coherence.

## Introduction

The molecular composition and pore size of the interstitium can vary substantially between tissue-types, physiological and inflammatory states, posing both physical and biochemical challenges for migrating immune cells. Mesenchymal cells like fibroblasts usually employ destructive locomotion strategies. These cells engage in tight adhesive interactions with the environment and use acto-myosin forces to pull on and deform the interstitial matrix^1^. Whenever transient deformation is not sufficient, proteolytic enzymes can also digest a path for passage of the cell body^2,3^. Mesenchymal cells typically position the centrosome and most organelles in front of the nucleus, where the secretory machinery is ideally positioned to deliver adhesion molecules and proteases^4,5^. In contrast, amoeboid cells like leukocytes, which migrate up to two orders of magnitude faster, are more opportunistic^6^. Usually, they neither permanently remodel their environment, nor tightly adhere to it. While migrating, leukocytes typically position their nucleus forward, followed by centrosome and organelles^5,7^. This configuration allows them to utilize the nucleus as a gauge to probe their vicinity, select larger pores over smaller ones and thereby navigate along a path of least resistance^8^. Amoeboid and mesenchymal configurations are paradigmatic locomotion strategies that seemed mainly determined by cell-intrinsic properties^9^. However, new evidence suggests that in response to specific environmental parameters like extreme confinement, inability to proteolyse and lack of adhesive ligands, mesenchymal cells can also adopt amoeboid features, a process termed mesenchymal to amoeboid transition^10,11^. To what extent the reverse is true and amoeboid cells can undergo an amoeboid to mesenchymal transition is less clear^12,13^.

Although amoeboid and mesenchymal cells operate in quantitatively very different force-regimes, they use the same force generating principles. These are mainly based in the actin cytoskeleton and can roughly be categorized as pulling or pushing forces. Pulling forces strictly require coupling to the environment via substrate-specific adhesion receptors. Thanks to a plethora of highly sophisticated traction force microscopy techniques, there is good quantitative understanding of how isolated adhesion sites locally pull on their substrate^14^. Less is known about pushing forces. These can result from cortical acto-myosin contractility due to the build-up of hydrostatic pressure, as exemplified in cellular blebs^15^. Alternatively, actin can also directly polymerize against and thereby protrude the plasma membrane as seen in lamellipodia and filopodia. To what extent pushing forces generated via actin polymerization are sufficient to locally displace or deform external obstacles like the extracellular matrix or neighboring cells is not firmly established^16^, but they seem especially relevant for amoeboid cells that do not transmit strong pulling forces via transmembrane adhesion receptors.

To ultimately understand how a cell translates intracellular forces into locomotion of the whole cell body it is important to not only study how one localized adhesion pulls or protrusion pushes on a substrate, but also how mechanical forces are coordinated on the scale of the whole cell. In order to address these aspects, we investigated how the cytoskeleton of amoeboid migrating Dendritic Cells (DCs) mechanically responds to controlled environments of high density and how this response affects overall cell shape dynamics.

## Results

### Dendritic cells undergo environmentally induced transition from amoeboid to mesenchymal configuration

To establish whether leukocytes resort to an alternative locomotion strategy in different environments, we observed the migration of mature DCs exposed to a gradient of the chemokine CCL19 in collagen gels of varying concentrations (1.7-3.5 mg/mL) **(Fig. 1a)**. In this setting, the cell population uniformly migrated in the direction of the chemokine gradient. After fixation, we quantified the relative position of nucleus and centrosome (eGFP-Centrin) along the polarisation axis of the cells. While in low density gels DCs predominantly migrated nucleus first, this orientation was reversed in high density gels **(Fig. 1b,c)**, indicating that DCs can switch from an amoeboid to a mesenchymal configuration when encountering narrow pores. To challenge this finding in controlled geometries we chemotactically guided EB3-*mCherry-*expressing DCs, which report microtubule plus end dynamics, through 1D microfluidic channels with narrow constrictions at the entrance **(Fig. 1d)**. Microtubule plus-end dynamics showed that virtually all microtubules originated from a single location, confirming that in DCs the centrosome serves as the sole microtubule organizing centre (MTOC). Like in collagen gels, organelle orientation was dependent on the cross section of the constriction, with the MTOC-first orientation being more prevalent in smaller cross-sections. When advancing through the straight, unconstricted part of the channel, cells frequently reverted to a nucleus first configuration **(Fig. 1e and Supplementary Video 1)**. Upon entering constrictions, cells co-expressing the actin reporter LifeAct-eGFP consistently showed an intense actin signal inside the constricted area that, like the MTOC, located in front of the nucleus **(Fig. 1d, f)**. The intensity of this actin signal increased with decreasing cross-section of the constriction **(Fig. 1g, h)**. We observed no obvious actin accumulation in cells migrating through straight channels **(Fig. 1d)**.

**Figure 1.**
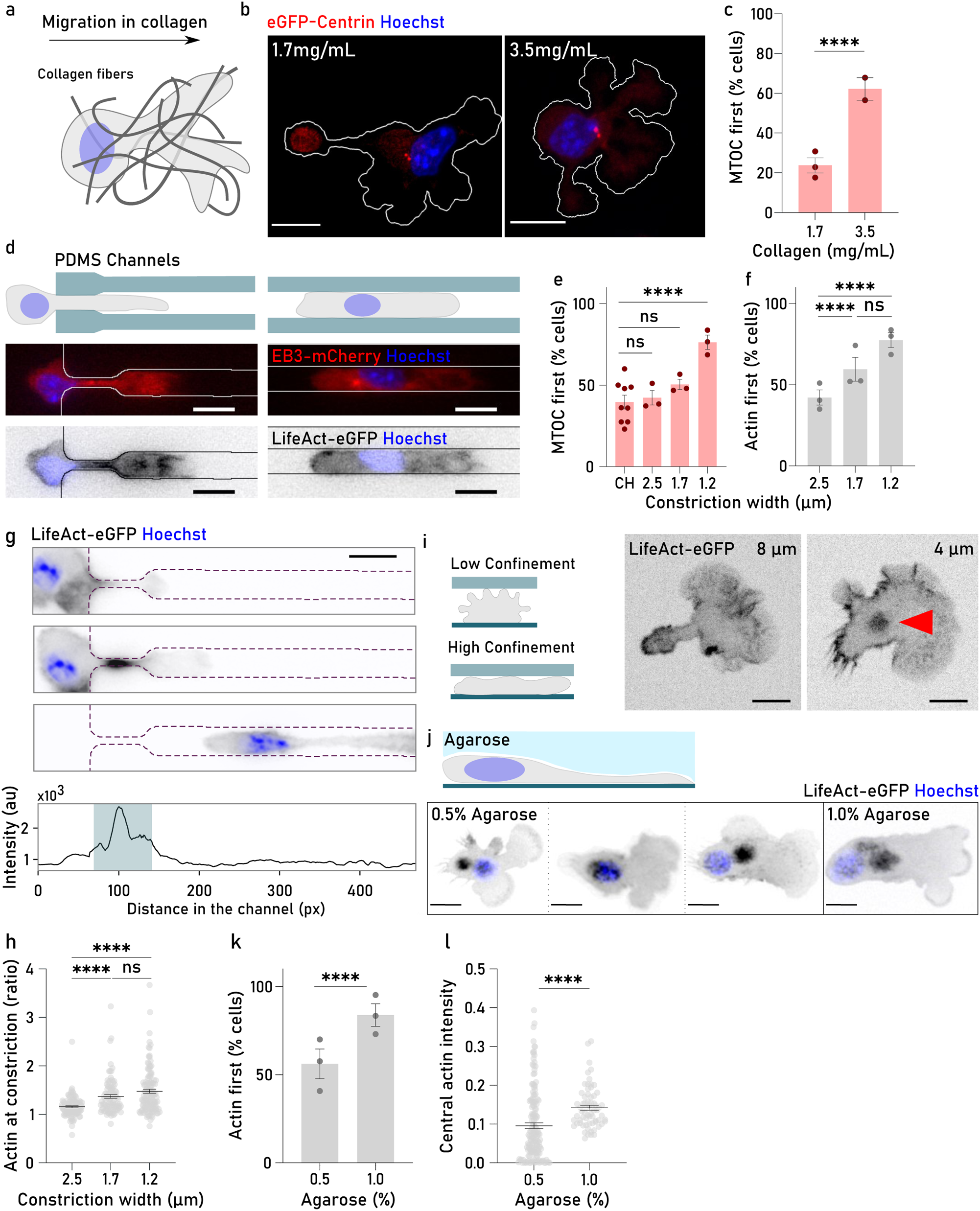
**a.** Schematic of a dendritic cell migrating in a collagen matrix. **b.** Representative images of dendritic cells moving in different concentrations of collagen (1.7 and 3.5 mg/mL). DCs expressed eGFP-Centrin (red) and were labeled with Hoechst (blue) for MTOC and nucleus visualization, respectively. Cell contour is shown in white. Scale bar, 10 µm. **c.** Percentage of cells migrating with MTOC first orientation in different collagen concentrations. Each dot represents the percentage of cells showing MTOC-first orientation in at least two independent experiments. Cells migrating in 1.7 mg/mL collagen, n=125; cells migrating in 3.5 mg/mL collagen, n=97. Error bars show the Standard Error of the Mean (SEM). **** p-value<0.0001. Fisher’s exact test. **d.** Migration in PDMS microchannels with narrow constrictions (1.2-2.5 μm width, 6 μm height), simulating transition through narrow pores (left), and migration in simple, straight PDMS microchannels (6 μm width, 6 μm height) mimicking cells migrating through large pores (right). EB3-*mcherry* (red) and LifeAct-eGFP (black) expressing DCS were used for MTOC and actin visualization, respectively. Hoechst was used for nuclear labeling (blue). Scale bar, 10 μm. **e.** Cells showing MTOC-first orientation during migration in straight channels (CH) or in constrictions of the indicated width. Each dot represents the percentage of cells showing MTOC-first orientation in at least three independent experiments. CH n=426; 2.5 μm n=137; 1.7 μm n=117, and 1.2 μm n=172. Error bars show the SEM. ns p-value=0.5495 (CH vs 2.5 μm), ns p-value= 0.0566 (CH vs 1.7 μm), and **** p-value<0.0001 (CH vs 1.2 μm). Fisher’s exact test. **f.** Cells showing actin-first orientation during migration in constrictions of the indicated width. Each dot represents the percentage of cells showing actin-first orientation in at least three independent experiments. 2.5 μm n=138; 1.7 μm n=119, and 1.2 μm n=162. Error bars show the SEM. **** p-value<0.0001, ns p-value=0.6405. Fisher’s exact test. **g.** Time lapse of a LifeAct-eGFP expressing cell (actin, black) entering a constricted channel (1.7 µm width). Note that actin accumulation in the constriction precedes the nucleus (Hoechst, blue). Scale bar, 10 μm. Bottom plot illustrates the temporal projection of maximum actin density of this cell moving in the constricted channel. Constriction is highlighted in blue. **h.** Ratio between maximum actin density at the constriction and outside of the constriction. Each dot corresponds to one single cell. Data was pooled from three independent experiments. 2.5 µm n= 101, 1.7 μm n=95, and 1.2 μm n=129. Error bars show the SEM. **** p-value<0.0001, ns p-value=0.5831. Unpaired Mann-Whitney test. **i.** LifeAct-eGFP (actin, black) expressing dendritic cells migrating in vertical PDMS confiners of different heights (left: 8 μm, right: 4 μm). Red arrow shows the actin accumulation in the middle of the cell in higher confinement. Scale bar, 10 μm. **j.** Cells migrating under agarose of different stiffness (0.5% and 1%). LifeAct-eGFP (actin, black) expressing DCs labeled with Hoechst (nucleus, blue). Note the different positioning of the central actin pool in relation to the nucleus in 0.5% agarose. Scale bar, 15 μm. **k.** Cells showing actin-first orientation when migrating under 0.5% or 1% agarose. Each dot corresponds to the percentage of cells where actin-first orientation was observed in three independent experiments. Cells migrating under 0.5% agarose, n=145; cells migrating under 1% agarose, n=88. Error bars show SEM. **** p-value<0.0001. Fisher’s exact test. **l.** Normalized intensity of the central actin in cells migrating under 0.5% or 1.0% agarose. Each dot shows mean intensity observed at the central actin pool, normalized to the global F-actin intensity of the cell over time. Data was pooled from three independent experiments. Cells migration under 0.5% agarose, n= 155; cells migrating under 1.0% agarose, n= 67. Error bars show SEM. **** p-value<0.0001. Unpaired Mann-Whitney test.

Actin accumulation in the constricted part of the channel suggested a response to compression of the cell body. We therefore vertically confined DCs between two surfaces separated by varying distances and imaged their LifeAct-eGFP distribution during chemokine driven locomotion. In this setting, cells showed a circular-shaped pool of actin that located in the cell center **(Fig. 1i and Supplementary Video 1)**. This central pool of actin was clearly distinguishable from both leading and trailing edge actin and constituted the most prominent actin pool in the DCs. In agreement with our findings, the number of cells showing the central actin pool increased when decreasing the confinement height **(Extended Data Fig. 1a)**.

Being confined within stiff environments (like in the microfluidic setting) and soft environments (like in vivo tissues, in collagen gels or under layers of soft material), can have different effects on migrating cells. We therefore imaged DCs migrating under agarose of varying concentrations. Here, completely confined cells have to lift the deformable layer of agarose in order to move **(Fig. 1j).** In this set-up, the central actin pool was present in virtually all cells migrating under both soft (0.5%) and stiff (1%) agarose **(Extended Data Fig. 1b)**. Under soft agarose only 50% of cells showed the central actin pool in front of the nucleus. The prevalence of this configuration increased to 80-85% under stiff agarose (1.0%) which was accompanied by a higher intensity of the central actin pool **(Fig. 1j-l and Supplementary Video 1)**. Similarly, we also observed a higher prevalence of MTOC-first migrating DCs in stiffer agarose **(Extended Data Fig. 1c, d and Supplementary Video 1)**. Other organelles like the Golgi apparatus and lysosomes, also positioned in front of the nucleus **(Extended Data Fig. 1e)**.

In summary, our data show that, whenever DCs have to transit through narrow spaces that demand extensive substrate and/or cell body deformations, they adopt a mesenchymal configuration. Associated with the MTOC and organelles, cells assemble a mechanosensitive central pool of actin that responds to different degrees of physical confinement.

### The central actin pool induces substrate deformations

Mechanoresponsiveness of the central actin pool suggested that one of its roles might be to counter external forces acting orthogonal to the direction of migration. To test this possibility, we developed pushing force microscopy, allowing us to associate substrate deformations with cellular structures. We let DCs migrate under agarose mixed with fluorescent beads, which we tracked in height with sub-micrometre precision using kymographic analysis of fast confocal microscopy stacks **(Fig. 2a, Extended Data Fig. 2a and Supplementary Video 2)**. Beads were stably incorporated into the agarose and remained stationary over time in the absence of cells. In contrast, beads were vertically displaced when cells migrated below them, indicating that cells could push against and transiently deform the agarose **(Fig. 2b)**. In order to locate more precisely which parts of the cell contributed to these deformations, we simultaneously imaged the nucleus and actin while probing bead displacement **(Fig. 2c)**. While beads were detectably displaced by the whole cell body, including the periphery, the displacement was most prominent right above the central pool of actin. Passage of the nucleus sustained the deformations induced by the central actin pool before substrate relaxation to its original position during nuclear exit. **(Fig. 2d)**. Cross-correlation analysis between bead-displacement and each of the imaging signals confirmed this order of events: bead displacement was strongly correlated with LifeAct-eGFP signal, while nuclear signal showed a weaker and asymmetric correlation **(Fig. 2e)**. In agreement with these results, we detected similar local substrate deformations associated with actin bursts in DCs migrating through collagen I matrices **(Extended Data Fig. 2b-d and Supplementary Video 3)**. Spatial maps of collagen fiber deformation and actin intensity maxima showed that local maxima of fiber displacements were in close proximity to peaks in LifeAct-eGFP signal. **(Extended Data Fig. 2e, f)**. These findings supported the notion that cells encountering confined spaces resort to actin polymerization in order to locally deform the extracellular environment.

**Figure 2.**
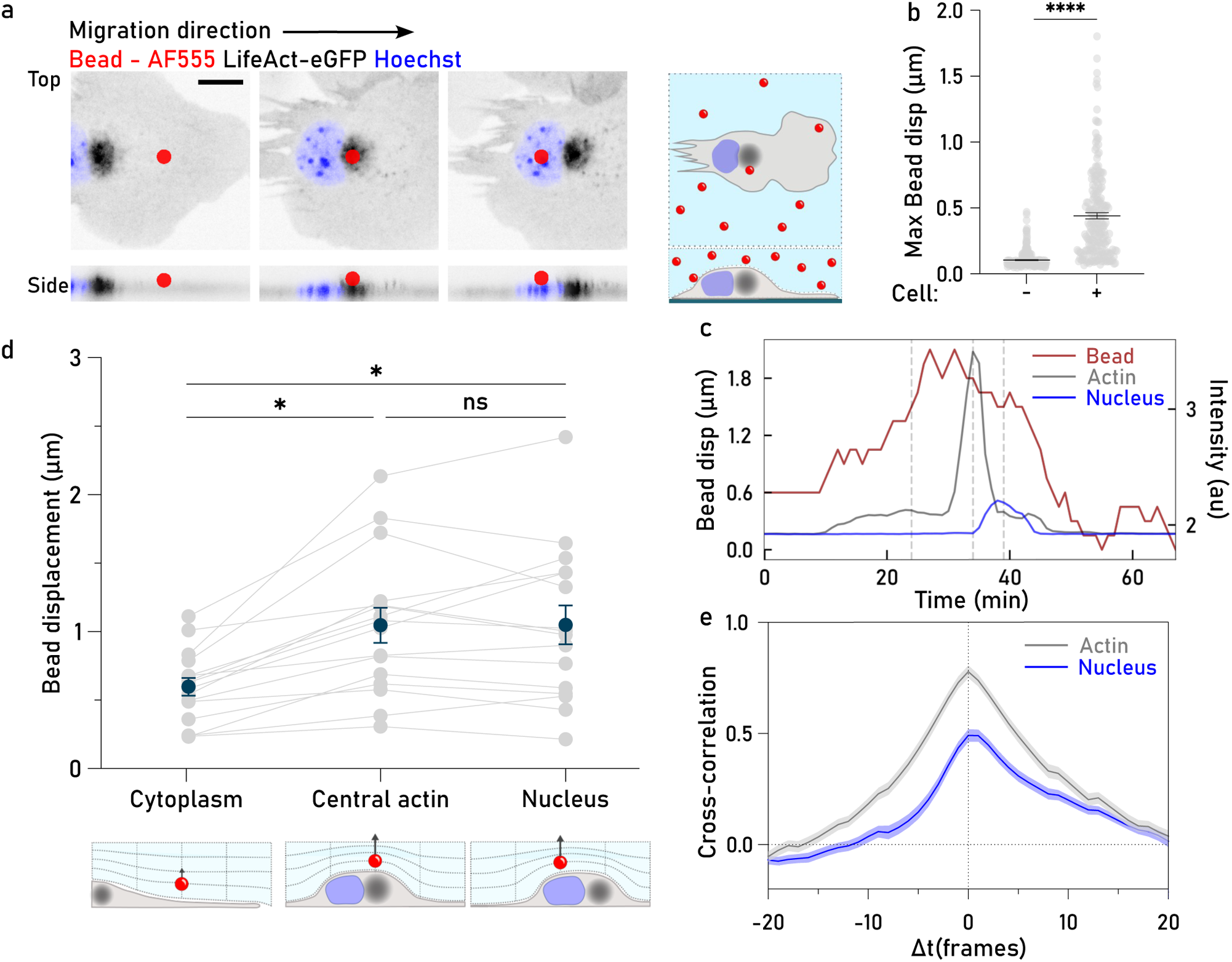
**a.** Cells migrating under agarose with fluorescent beads labeled with AF555. While the upper panel shows the top view, the lower panel shows the lateral projection of a migrating cell. Three different time points are represented: (1) cell body under the bead; (2) central actin cloud under the bead, and (3) nucleus under the bead. Cells expressed LifeAct-eGFP (actin, black) and were labeled with Hoechst (nucleus, blue). Scale bar 10 μm. **b.** Maximum bead displacement in the absence (−) or presence (+) of DCs migrating under the beads. Each dot corresponds to the displacement observed for an individual bead and cell. Bead displacement in the absence of cells, n=333; bead displacement in the presence of cells, n=205. Error bars show SEM. **** p-value<0.0001. Unpaired Mann-Whitney test. **c.** Change in bead displacement in Z (red, left) or LifeAct-eGFP (actin, black), and Hoechst (nucleus, blue) intensities over time. The dashed gray lines highlight the three time points shown in A. **d.** Cytoplasmic, central actin pool, and nuclear contribution for bead displacement in Z. Each dot corresponds to the displacement of one single bead from at least three independent experiments, n=16. A line connects measurements corresponding to the same bead. Dark blue dots show the mean displacement in Z and error bars show SEM. * p-value=0.0241 (cytoplasm vs central actin), * p-value=0.0230 (cytoplasm vs nucleus), and ns p-value=0.9998 (central actin vs nucleus). One-way ANOVA. **e.** Temporal cross-correlation between bead displacement and nucleus intensity (blue) or actin intensity (grey). Shadow error bars show SEM.

### Cdc42 and its exchange factor DOCK8 regulate the central actin pool

Next, we wondered how perturbations of the central actin pool would impact the ability of cells to migrate and interact with the substrate. Actin polymerization in leukocytes is predominantly controlled by two canonical pathways: the small Rho GTPases Rac1 and Cdc42. Rac1 mainly triggers actin polymerisation via direct interaction with the WAVE complex, which in turn activates Arp2/3 dependent nucleation of new branched filaments. While WAVE locates at the tip of lamellipodia and is essential for the typical veiled shape of DCs^17,18^, Cdc42 has more pleiotropic effects on cytoskeletal dynamics and cell polarity^19^. Among other effectors, Cdc42 triggers WASP dependent Arp2/3 activation, which has complex and still poorly-understood cellular functions, ranging from local, curvature-dependent force generation^20,21^ to the formation of invadosomes^22^ and endocytic sites^23^. To probe whether and how these pathways affected the central actin pool while avoiding deranging the homeostasis of cell shape and membrane dynamics, we treated DCs migrating under agarose with low concentrations of Rac1 and Cdc42 inhibitors **(Extended Data Fig. 3a, b and Supplementary Video 4)**. While we could not detect any obvious effects by Rac1 inhibition **(Extended Data Fig. 3c, d)**, Cdc42 inhibition led to a decrease in the prevalence of the central actin pool to only 50% of cells and a reduction of the local F-actin signal in the central pool **(Fig. 3a-c, Extended Data Fig. 3e)**. Interestingly, no changes in the total amount of cellular F-actin were observed **(Fig. 3d)**. Transient transfection of DCs with dominant negative GFP-Cdc42(T17N) confirmed the key regulatory role of Cdc42 for the central pool of actin **(Fig. 3e, f)**.

**Figure 3.**
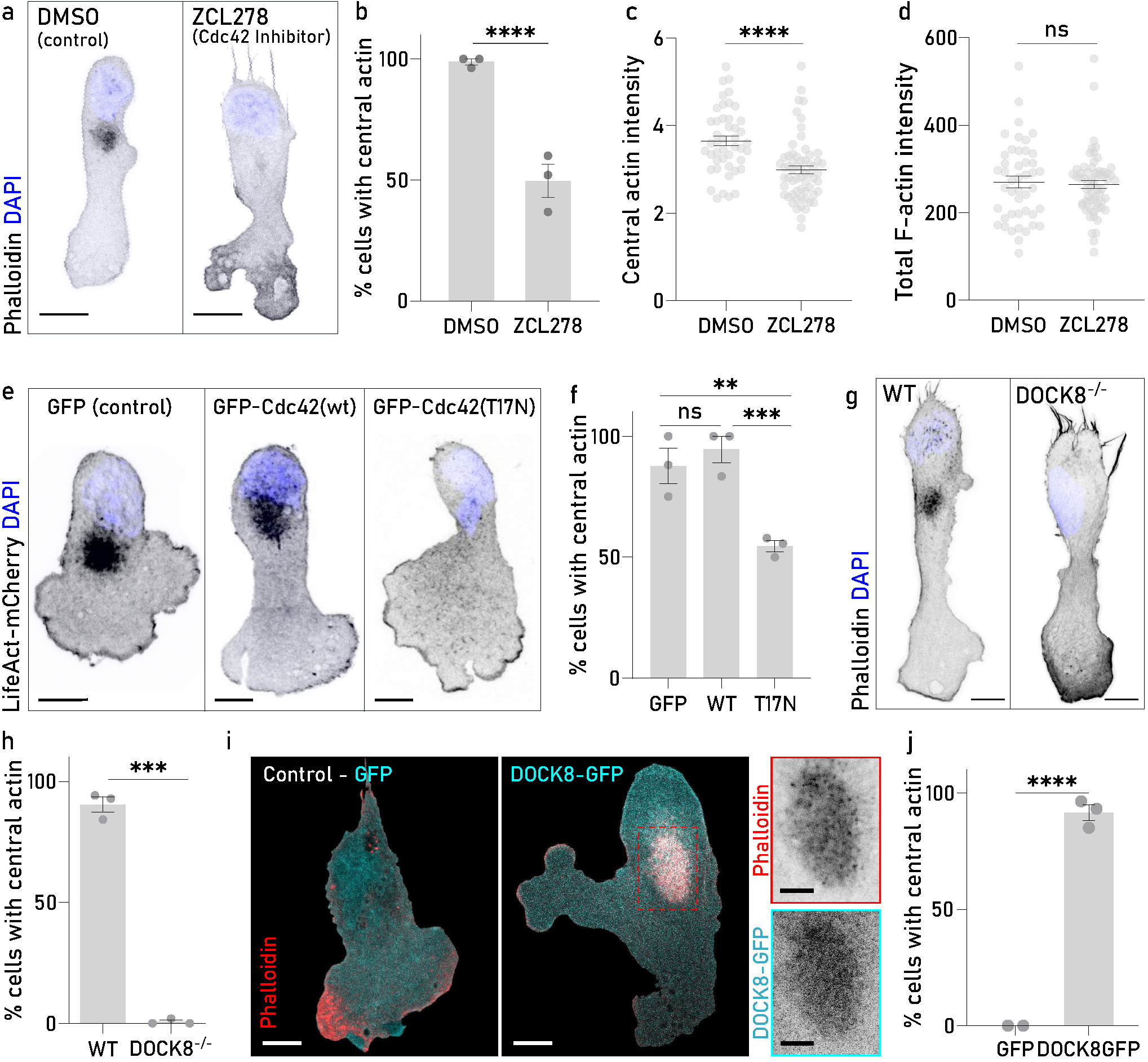
**a.** DCs migrating under a patch of 1% agarose treated either with DMSO (control - left) or ZCL278 (Cdc42 inhibitor - right). Cells were fixed and stained with phalloidin (F-actin, black) and DAPI (nucleus, blue). Scale bar, 10 μm. **b.** Cells showing a central pool of actin upon treatment with DMSO or ZCL278. Each dot represents the percentage of cells where a central actin pool was detectable in three independent experiments. DMSO treated cells, n= 147; ZCL278 treated cells, n=66. Error bars show the SEM. **** p-value<0.0001. Fisher’s exact test. **c.** Central actin intensity of cells treated with ZCL278 or control cells (DMSO). Each dot corresponds to the mean central actin intensity normalized to the global actin intensity in one individual cell. Data was pooled from three independent experiments. DMSO n=45; ZCL278 n=61. Error bars show SEM. **** p-value<0.0001. Unpaired Mann-Whitney test. **d.** Total F-actin intensity measured in DMSO or ZCL278 inhibited DCs. Each dot corresponds to the mean F-actin intensity in one individual cell. Data was pooled from three independent experiments. DMSO n=45; ZCL278 n=61. Error bars show SEM. ns p-value=0.7455. Unpaired Mann-Whitney test. **e.** DCs migrating under a patch of 1% agarose after transfection with one of the following constructs: GFP (control), GFP-(wt)Cdc42 (WT), or GFP-(T17N)Cdc42 (T17N). Positive construct transfection was confirmed by GFP expression. Cells were stained with phalloidin (F-actin, black) and DAPI (nucleus, blue). Scale bar, 10 μm. **f.** Cells showing central actin after transient transfection with the indicated constructs. Each dot corresponds to the percentage of cells with a detectable central actin pool observed in three independent experiments. GFP, n= 32; WT, n=27; T17N, n=52. ns p-value =0.6780, ** p-value = 0.0018, *** p-value = 0.0004. Fisher’s exact test. **g.** WT and DOCK8^−/−^ fixed DCs migrating under a patch of 1% agarose. Cells were fixed and stained with phalloidin (F-actin, black) and DAPI (nucleus, blue). Scale bar, 10 μm. **h.** Central actin presence in WT and DOCK8^−/−^ DCs migrating under agarose. Each dot corresponds to the percentage of cells showing a detectable central actin pool for three independent experiments. WT, n= 240; DOCK8^−/−^, n= 171. **** p-value<0.0001. Fisher’s exact test. **i.** DOCK8^−/−^ DCs expressing GFP (cyan, left) or WT-DOCK8-GFP (cyan, right). Cells were fixed and stained with phalloidin (F-actin, red). Scale bar, 10μm. The red dashed box marks the area used for the inset. Top: Phalloidin (F-actin, red) showing the rescue of the central actin pool; Bottom: WT-DOCK8-GFP (DOCK8, cyan) showing DOCK8 localization in the central actin region. Scale bar, 5 μm. **j.** Central actin in DOCK8^−/−^ DCs expressing GFP or WT-DOCK8-GFP migrating under agarose. Each dot corresponds to the percentage of cells showing a detectable central actin pool for at least two independent experiments. GFP, n=47; WT-DOCK8-GFP, n=92., **** p-value<0.0001. Fisher’s exact test.

Cdc42 interacts with several guanine exchange factors (GEFs). Among them DOCK8, which is prominently expressed in the hematopoietic lineage and causative for a severe congenital immunodeficiency associated with actin dysregulation^24–26^. In line with previous studies, cells with a genetic deletion of DOCK8 (DOCK8^−/−^) showed regular differentiation and their typical veiled morphology was indistinguishable from wild type (WT) cells when floating in suspension^27^. Strikingly however, when these cells were confined under agarose they showed a complete lack of the central actin pool **(Fig. 3g, h)**. Re-expression of DOCK8-GFP in DOCK8 deficient cells was sufficient to rescue the WT phenotype, and revealed that DOCK8 colocalized with the central actin pool and was not present anywhere else throughout the cell, including the leading edge **(Fig. 3i, j)**. Together, these data show that DOCK8 is the key regulator of the mechanosensitive central pool of actin.

### Central actin communicates with leading edge actin

To investigate the role of the central pool of actin in cell motility, we imaged the chemotactic migration of DOCK8^−/−^ DCs confined under agarose. The migration speed of DOCK8^−/−^ DCs was not different compared to control cells **(Fig. 4a)**. However, DOCK8^−/−^ cells inflicted smaller actin-mediated deformations on the agarose **(Fig. 4b)**, with the nucleus now being the main bearer of the load in the absence of the central actin pool **(Fig. 4c)**. In addition, DOCK8^−/−^ DCs displayed distinct elongated morphology and an incoherent leading edge that often branched in two or more lobes **(Fig. 4d and Supplementary Video 5)**. In confined environments, lamellipodial splitting is a typical sign of stabilization, as opposed to destabilized lamellipodia that show more coherent morphology. Indeed, phalloidin staining of fixed cells revealed that, while global F-actin levels were minimally reduced compared to control cells **(Extended Data Fig.4 a, b)**, F-actin signal was substantially enhanced at the leading edge of DOCK8^−/−^ DCs (**Fig. 3g**, **Fig. 4e, Extended Data Fig. 4c**). Together, these data suggest that the lack of the central actin pool in DOCK8^−/−^ DCs triggers a new locomotory equilibrium. The inability to deform the substrate results in jamming of the cell body, which is compensated by an hyperstabilized leading edge which drags the cell forward.

**Figure 4.**
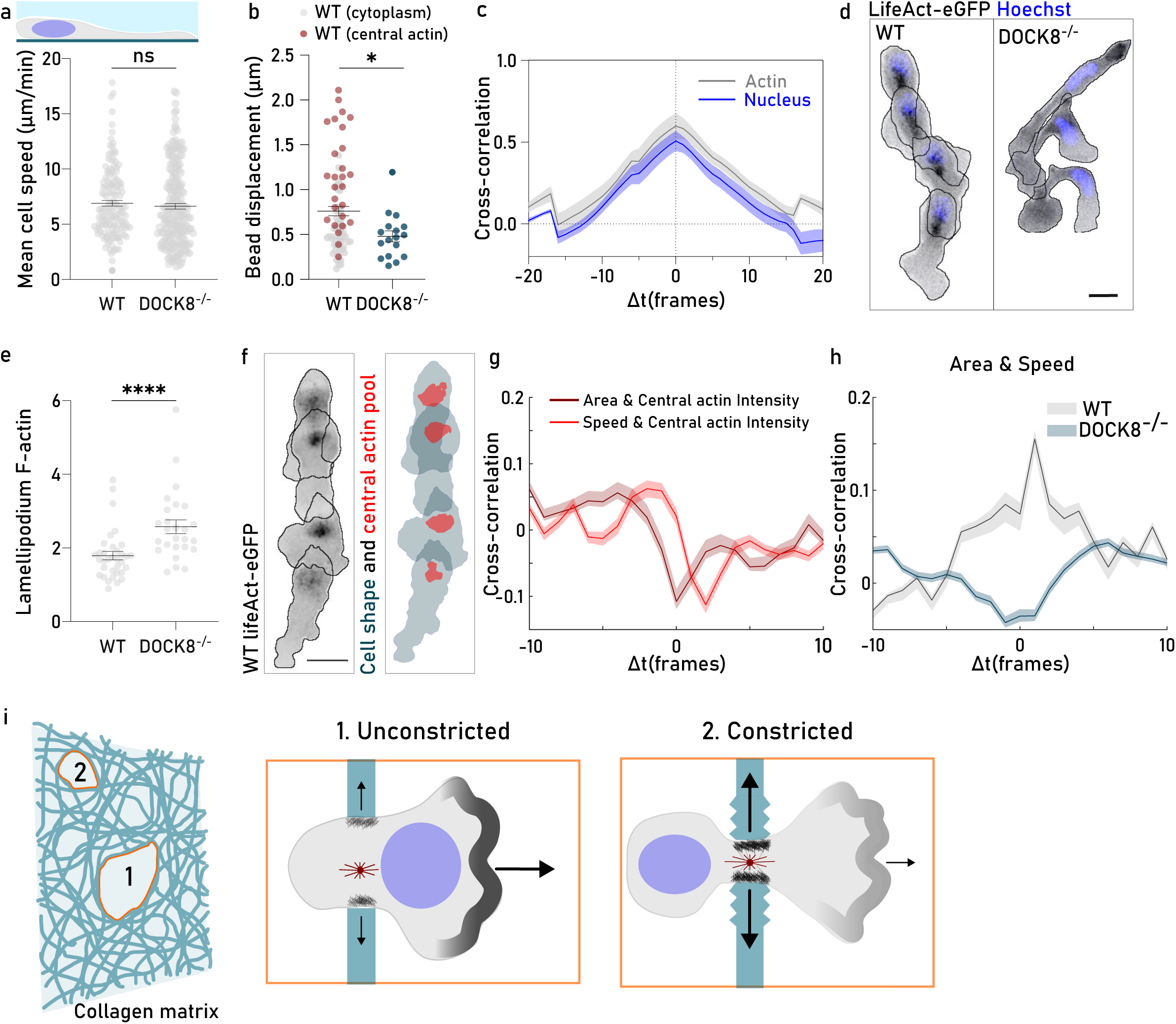
**a.** WT and DOCK8^−/−^ cells migration under a patch of 1% agarose. Each dot corresponds to the mean speed observed for one single cell over time. Data was pooled from at least three independent experiments. WT, n=86; DOCK8^−/−^, n=111. Error bars show SEM. ns p-value=0.1658. Mann-Whitney test. **b.** Bead displacement generated either WT or DOCK8-/- DCs. Each dot corresponds to a single bead displaced by the cell body (excluding the nucleus). To avoid bias, only beads under which the nucleus passed were considered. Within WT DCs, red dots represent beads that were also displaced by the central pool of actin, reflecting larger deformations. WT, n=80; DOCK8-/-, n=17. Error bars show SEM. * p-value=0.0239. Mann-Whitney test. **c.** Temporal cross-correlation between bead displacement generated by DOCK8-/- DCs and nucleus intensity (blue) or actin intensity (grey). Shadow error bars show SEM. **d.** Time-lapse projection of WT (left) and DOCK8^−/−^ (right) DCs expressing LifeAct-eGFP (F-actin, black) and labeled with Hoechst (nucleus, blue) migrating under 1% agarose. Scale bar, 15 μm. **e.** F-actin density at the lamellipodium of WT and DOCK8^−/−^ cells migrating under agarose. Each dot corresponds to the mean F-actin density at the lamellipodium normalized to the total F-actin density for one individual cell. WT, n=34; DOCK8^−/−^, n=26. Error bar show SEM. **** p-value<0.0001. Mann-Whitney test. **f.** Left: Time-lapse projection of a migrating WT LifeAct-eGFP (actin, black) expressing DC. Contour of the cell is shown by the black line. Scale bar, 15 µm. Right: Cell (blue) and central actin pool (red) segmentation results. Segmented areas were manually curated and used for the subsequent quantifications. **g.** Temporal cross-correlation between central actin intensity and cell speed (red), or cell area (dark red) in WT DCs migrating under 1% agarose. Data points were pooled from three independent experiments, n=68. Shadow error bars show SEM. **h.** Temporal cross-correlation between cell area and cell speed in WT (light blue) and DOCK8^−/−^ (dark blue) DCs migrating under 1% agarose. Data points were pooled from three independent experiments. WT n=68, DOCK8^−/−^ n=177. Shadow error bars show SEM. **i.** The surrounding environment of a cell dictates organelle orientation and triggers changes in the actin cytoskeleton. In unconstricted environments migrating DCs show a nucleus-first configuration and an actin enriched leading edge (1). When cells encounter constrictions, actin is recruited to the central pool (represented by the black hashing). Increased actin polymerization at the central pool promotes deformation of the surrounding environment and results in a reduction of actin at the leading edge (2).

To better understand the communication between actin at the cell front and actin at the cell body, we turned to WT cells and quantitatively imaged the dynamics of the central pool of actin, cell shape and locomotion. We first tested WT cells in PDMS pillar mazes where small obstacles in the migratory path promote splitting of the lamellipodium **(Extended Data Fig. 4d)**. We observed that lamellipodium retractions were accompanied by an increase of actin intensity at the central pool (**Extended Data Fig. 4e, f and Supplementary Video 5**), while no significant actin signal variations were detected in other areas of the cell body (**Extended Data Fig. 4f**). This observation was corroborated in cells migrating under agarose **(Fig. 4f and Supplementary Video 5)**. Here, cell speed and central actin intensity showed a strong negative correlation **(Fig. 4g)**, suggesting that reductions of the central actin pool are associated with a “run phase”, while stalled migration corresponds to an increase in the central actin pool. The intensity of the central actin pool and the projected cell area also showed a slightly delayed, negative correlation **(Fig. 4g)**, reflecting cell spreading whenever the central actin pool shrinks. Consistent with these patterns, cell speed was positively correlated with the projected area in WT cells. This correlation was lost in DOCK8^−/−^ DCs **(Fig. 4h)**.

From our data, we propose a model in which cells redistribute actin between the leading edge and a central pool of actin on demand **(Fig. 4i)**. Under favorable migratory conditions in which the cell body is largely unobstructed (i.e. in the absence of tight constrictions), actin is enriched at the cell front, enhancing leading edge protrusion and accelerating forward locomotion. In contrast, when cells face more constrictive environments, actin is recruited to the central pool. This solution offers a double advantage: on the one hand, actin polymerisation dilates a path for organelles and nucleus, preventing cell entrapment in areas of high confinement. At the same time, it serves as a “capacitor” for actin by restricting actin accumulation at the cell front, thereby preventing leading edge advancement whenever the cell body is trapped.

### DOCK8 differentially affects DC locomotion depending on environmental factors

Our model implies that dysregulation of the central actin pool can have a different impact on cell migration depending on the geometry and complexity of the environment. In line with previous findings^27–29^, DOCK8^−/−^ cells migrating in collagen gels were substantially slower than WT cells and showed signs of enhanced leading edge stabilization **(Fig. 5a, b and Supplementary Video 6)**. Their phenotype of extending multiple simultaneous protrusions was even more pronounced than under agarose **(Fig. 5c).** Moreover, and consistent with findings in T cells^30^, we observed a high rate of fragmentation in DOCK8 deficient DCs, which often resulted in cell death **(Fig. 5d, e)**. Interestingly, the resulting cell fragments, especially those originating from the leading edge, were often motile and chemotactic **(Fig. 5f).** DOCK8^−/−^ DCs chemotactically migrating in relatively complex PDMS pillar mazes (with 1-μm, 2-μm or 3-μm-distanced pillars) reproduced the phenotype observed in collagen gels: cells extended multiple protrusions, entangled and often fragmented **(Fig. 5g and Supplementary Video 6)**. However, occasional DOCK8^−/−^ DCs that adopted a monopolar configuration migrated significantly faster than cells with multiple competing leading edges **(Fig. 5h).** These observations are consistent with our model **(Fig. 4i)**. Lack of the central actin pool not only leads to actin accumulation at the leading front of DOCK8^−/−^ DCs promoting its advancement, but also precludes substrate deformations by the cell body. This results in entanglement and fragmentation of DOCK8^−/−^ DCs when migrating in complex fibrillar meshworks.

**Figure 5.**
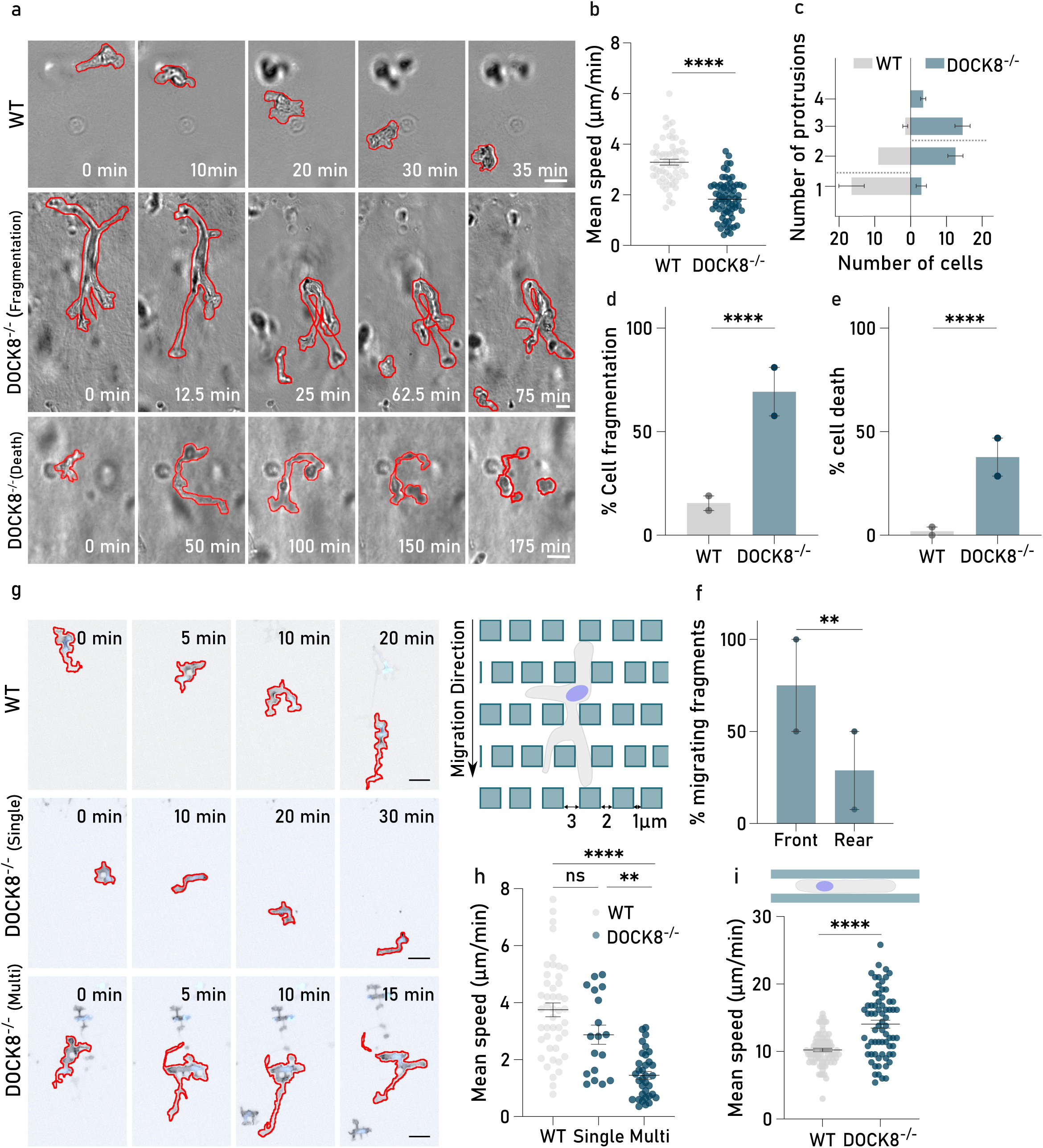
**a.** Brightfield images of WT (top) and DOCK8^−/−^ (middle, bottom) DCs migrating in 1.7 mg/mL collagen. While the middle panel shows an example of DOCK8^−/−^ cell fragmenting, the bottom panel shows a DOCK8^−/−^ DC undergoing apoptosis during migration. Contour of the cells is shown in red. Scale bar, 15 μm. **b.** WT (grey) and DOCK8^−/−^ (blue) cells migration in 1.7 mg/mL collagen. Each dots represents the mean speed of an individual cell over time. Data was pooled from two independent experiments. WT, n=59; DOCK8^−/−^, n=69. Error bar show SEM. **** p-value<0.0001. Unpaired t-test. **c.** Number of simultaneous protrusions observed in one individual cell in WT (left) and DOCK8^−/−^ (right) DCs during migration in 1.7 mg/mL collagen matrices. Data was pooled from two independent experiments. Dashed lines correspond to the mean number of protrusions observed during migration and error bars show SEM. WT, n=54; DOCK8^−/−^, n= 67. (p-value<0.0001. Fisher’s exact test.) **d.** Fragmentation rate of WT (grey) and DOCK8^−/−^ (blue) cells migrating in a 1.7mg/mL collagen matrix. Each dot corresponds to the percentage of cells where we detected fragmentation events in two independent experiments. WT, n=54; DOCK8^−/−^, n= 67. Error bars show SEM. **** p-value<0.0001. Fisher’s exact test **e.** Cell death rate during migration in 1.7mg/mL collagen. WT is shown in grey; DOCK8^−/−^ is shown in blue. Each dot corresponds to the percentage of cells that underwent apoptosis in two independent experiments. WT, n=54; Dock8^−/−^, n=67. Error bars show SEM. **** p-value<0.0001. Fisher’s exact test. **f.** Percentage of migrating fragments originated from DOCK8^−/−^ cells. Front refers to fragments that originated from a protrusion, while rear indicates fragments originated from the rear end of a cell. Each dot to the percentage of these events observed in 2 independent experiments. Front n=11; Rear n= 25. ** p-value=0.0042. Fisher’s exact test. **g.** DCs moving in a constrictive pillar maze, confined between surfaces 6 µm apart and intersected by 1-µm, 2-µm or 3-µm-distanced pillars. Top: WT; middle and bottom: DOCK8^−/−^ DCs. DOCK8^−/−^ cells with a monopolar configuration (one single coherent lamellipodium – middle panel) migrate more efficiently in comparison with a cells with multiple lamellipodia (bottom panel). Contour of the cells is shown in red. Scale bar, 20 μm. **h.** Migration speed of WT, DOCK8^−/−^ cells with a single lamellipodium (Single) and DOCK8^−/−^ cells with multiple lamellipodia (Multi). Each dot corresponds to the mean speed of an individual cell through time. Data was pooled from three independent experiments. WT n=45, DOCK8^−/−^ single n=18, and DOCK8^−/−^ multi n=34. Error bars show SEM. **** p-value<0.0001, ** p-value=0.0033, ns p-value=0.2748. Mann-Whitney test. **i.** WT and DOCK8^−/−^ migration speed in straight PDMS channels. Each dot corresponds to the mean speed observed for an individual cell through time. Data was pooled from three independent experiments. Error bars show SEM. WT, n=83; DOCK8^−/−^, n= 69. **** p-value<0.0001. Unpaired t-test.

The model also predicts that enhanced front protrusions should boost forward locomotion in simple geometries where confinement and leading edge splitting is limited. We therefore tested DCs migrating in straight microfluidic channels with 6 µm diameter. In this set-up we found that DOCK8^−/−^ DCs were morphologically indistinguishable from WT cells and, more importantly, they now migrated substantially faster than WT cells **(Fig. 5i and Supplementary Video 6)**. Thus, if cells are not slowed down by restrictions imposed by the environment, redistribution of the central actin pool towards the leading edge enhances migration.

## Discussion

To move as a coherent unit, different parts of the cell have to act in coordination. For a leukocyte in migration-permissive environments, front to back polarity ensures the necessary coordination between protrusion and retraction. This coordination strongly depends on posterior actomyosin-contraction, which not only retracts the trailing edge but also squeezes the cell body and propels the nucleus to the front, locating it directly behind the lamellipodium^31–33^. In this configuration, cells use the bulky nucleus to choose larger pores over smaller ones^8^. Whenever the cell faces a situation where even the biggest pore does not allow passage, it can either stall, revert or create a new path. In increasingly dense and complex environments, typical strategies are not sufficient, because the cell has to simultaneously exert forces in multiple directions^20,21^. Nuclear propulsion raises the possibility that the nucleus aids in path dilation by serving as a wedge to dilate small pores^34^. However, nuclear and genomic integrity is sensitive to large forces^35,36^, indicating that the nucleus should rather be protected from excessive deformation than used as a force transducer.

Here we show that, when confronted with very narrow constrictions, amoeboid cells can switch to another strategy: they change their polar configuration from amoeboid to mesenchymal, where the nucleus is swept to the back and the MTOC is positioned in front. The local confinement imposed by the environment also triggers polymerization of a central pool of actin that associates with the MTOC and the bulk of cellular organelles. Actin polymerization in this central region generates pushing forces that not only deform the surrounding environment of the cell, but can potentially protect the nucleus and other organelles from fatal damage. This central actin pool is controlled by the activity of the Cdc42 GEF DOCK8 through a mechanosensitive pathway that remains to be identified. Intriguingly, an upstream activator of DOCK8 is the Hippo kinase MST1^37^, raising the possibility that this key mechanosensitive pathway, that regulates organ shape and size via controlling cell proliferation, also controls single cell deformation via its non-canonical effectors DOCK8 and Cdc42.

How different actin pools communicate is poorly understood in animal cells^38^ but better studied in yeast, where F-actin either comes in the form of patches or cables. Here, reduction of one structure is balanced by increase of the other, leaving the overall levels of F actin conserved^39^. We find a similar homeostatic balance in DCs and describe a regulatory loop between the central and the leading edge pools of actin. Our results suggest that via this communication axis, cells can coordinate protrusions in two orthogonal directions. Accordingly, DOCK8 deficient cells that lack this coordination do fragment, because a chemotactically enhanced leading edge loses contact with an immobilized cell body that is unable to push obstacles away. Together, our findings establish a novel regulatory loop between cell front and cell body that is essential for maintaining cellular coherence.

## Methods

### Mouse strains

*C57BL/6 (Janvier); LifeAct-eGFP*^40^; *DOCK8^−/−^* ^27^ (a gift from Yoshinori Fukui’s lab).

All mice used in this study were bred on a *C57BL/6* background and maintained at the Institute of Science and Technology Austria Institutional animal facility following the guidelines from its ethics commission and the Austrian law for animal experimentation.

### Generation and maintenance of immortalized hematopoietic progenitor cells

Hematopoietic progenitor cells were generated from the isolated bone marrow of 8 to 10-week-old mice which were retrovirally infected with an estrogen-regulated form of HoxB8 as described before^41^. Conditionally immortalized early hematopoietic progenitor cells were kept in R10 medium (RMPI 1640 supplemented with 10% fetal calf serum (FCS), 2 mM L-glutamine, 100 U/mL penicillin, 100 μg/mL streptomycin, and 50 μM β-mercaptoethanol (all Invitrogen), supplemented with 0.01% β-estradiol and 5% of in-house-generated Flt3tl containing supernatant). All cells were kept at 37 °C and 5% CO_2_ until differentiation.

### Constructs used for reporter progenitor cell lines

eGFP-Centrin cells were generated from a human centrin1 construct (a gift from Ana-Maria Lennon-Dumenil’s lab). EMTB-*mCherry* and EB3-*mCherry* DCs were generated as described before^42^. GFP and DOCK8-constructs GFP^43^ (a gift from Yoshinori Fukui’s lab) were modified to a pLenti6.3 backbone using Gibson Assembly strategy.

### Lentivirus production and transduction into progenitor cells

Fusion-protein-coding lentiviruses were produced in Lenti-X-293 cells derived from HEK 293 cells (TakaraBio). Lenti-X-293 cells were maintained in DMEM (Invitrogen) at 37 °C and 5% CO_2_ and transfected with the above-mentioned plasmids and two helper plasmids in Optimem (Invitrogen) and PEI (1 mg/mL, Polysciences). The supernatant was collected 48 hours after transfection and the resulting lentivirus preparation was concentrated using Lenti-X^TM^ Concentrator (Clontech) according to the manufacturer’s instructions. Progenitor cells were transduced with the concentrated lentiviral preparations by spin infection (1500 g, 1 hour) with 8 μg/mL Polybrene. Cells expressing the virus insertion were sorted in a Sony SH800 SFP cell sorter (sorting chip: 100 µm) for *mCherry* or GFP expression before DC differentiation.

### Differentiation and maturation of dendritic cells

Dendritic cells (DCs) were differentiated by seeding 3×10^5^ precursor cells in a 10 mL dish containing R10 medium supplemented with 10% of in-house-generated granulocyte-macrophage colony-stimulating factor (GM-CSF) hybridoma supernatant. On the third day of differentiation, 10 mL of R10 medium containing 20% GM-CSF was added to each dish. Half of the medium was replaced with R10 medium containing 20% GM-CSF on day 6 and cells were either harvested for maturation or frozen on day 8. Maturation was induced by overnight stimulation with lipopolysaccharide (LPS) from *Escherichia coli* 0127:B8 (Sigma) at a final concentration of 200 ng/mL.

### Flow Cytometry analysis of DCs

DCs were routinely checked for correct surface expression markers using antiboides against MHCII and CD11c (48-5321-82 and 17-0114-82, respectively, both eBiosciences). Stainings were performed in FACS buffer (1xPBS, 2 mM EDTA, 1% BSA) with Fc receptor blockage (anti-mouse CD16/CD32). Analysis was carried either on a FACSCanto BD Biosciences or in a BC CytoFLEX LX.

### Transient transfection of dendritic cells

The following plasmids were used: eGFP, pcDNA3-EGFPCdc42(wt), and pcDNA3-EGFP-Cdc42(T17N)^44^ (gifts from Klaus Hahn to Addgene, plasmids #12599 and #12601). DCs derived from progenitor cells were transfected with 4 μg of DNA using the nucleofector kit for primary T cells (Amaxa, Lonza Group) following the manufacturer’s guidelines. Briefly, 4-5×10^6^ cells were resuspended in 100 μL of DMEM (Invitrogen) and 4 μg of plasmid DNA. Cells were transferred to a cuvette and electroporated using a specifically designed protocol (program X-001). Transfected DCs were incubated overnight in R10 supplemented with 10% GM-CSF and LPS (200 ng/mL). Experiments were carried out the next day, and only GFP-expressing cells were analyzed.

### Pharmacological Inhibitors

The following small molecule inhibitors were used: ZCL278, to perturb Cdc42 activity^45^ (MedChemExpress); NSC23766 to perturb Rac1 activity^46^ (MedChemExpress), Inhibitors were mixed with the DC suspension after maturation for at least 30 minutes and kept through the assays at the indicated final concentration. ZCL278, 10 μM; NSC23766, 50 μM.

### FACS F-actin analysis in DCs

After overnight stimulation with LPS, WT and DOCK8^−/−^ DCs were recovered in 12-well plates in 500 μL R10 for 30 minutes at 37 °C. Cells were stained during fixation (4% PFA, 20 μM FITC-phalloidin and 0.5% saponin in phosphate-buffered saline (PBS), 500 μL, 20 minutes at 37 °C) and analyzed on a FACS Aria III. Stainings were carried out in three biologically independent samples.

### Under-agarose migration assay of mature dendritic cells

Glass coverslips were glued to the bottom of a petri dish with a 17 mm diameter hole where a custom-made plastic ring was attached using paraffin (Paraplast X-tra; Sigma). Agarose solution was prepared by mixing one part of 2x Hank’s buffered salt solution (HBSS, Sigma), pH 7.3 with 2 parts RPMI (Invitrogen) supplemented with 20% FCS (Invitrogen) and 2x the concentration of all the other supplements used in R10 medium (see above) and either 2% or 4% of UltraPure Agarose (Invitrogen) dissolved in one part water to achieve different agarose stiffnesses. 400 μL of the liquid agarose was poured into the dish, covering the coverslip. The agarose was allowed to solidify at room temperature for 5 minutes, after which two holes (1.5 mm and 2.0 mm) were punched into the agarose. The dishes were incubated at 37 °C and 5% CO_2_ for 30 minutes for equilibration. 2.5 μg/mL of CCL19 (PrepoTech) diluted in R10 was placed in the 2 mm hole and 0.5-1×10^6^ mature DCs were placed in the 1.5 mm hole opposite to the chemokine. Before acquisition, dishes were incubated for at least 1 hour at 37°C and 5% CO_2_ to allow invasion under the agarose. All images were acquired under physiological conditions using custom-built climate chambers (37 °C, 5% CO_2_, humidified).

### Immunofluorescence under-agarose

For analysis of fixed samples, a round-shaped coverslip (#1.5, 10 mm, Mentzel, Thermo Fisher Scientific) was placed in a glass-bottom dish before casting the agarose. DCs and chemokine were added to the dishes as described above. Cells were allowed to invade and migrate for at least 3 hours at 37 °C and 5% CO_2_. Migrating cells were fixed with prewarmed PBS supplemented with 4% paraformaldehyde (PFA) for 20 minutes at 37 °C. After fixation, the agarose patch was carefully removed and the coverslip was recovered and thoroughly washed with PBS. Coverslips were incubated with 0.1% Triton X-100 in PBS for 20 minutes at room temperature, washed with PBS, and blocked with 1% Bovine Serum Albumin (BSA) for 1 hour at room temperature. Primary antibodies were diluted in PBS with 1%BSA and incubated either overnight at 4 °C or for 2 h at room temperature. After primary antibody incubation, cells were washed 3 times with PBS and incubated with secondary antibody diluted in PBS with 1% BSA 1 hour at room temperature. Stained coverslips were washed 3 times with PBS and mounted on a slide using Flourmount-G mounting medium with DAPI (00-4959-52, ThermoFisher Scientific). Slides were imaged the next day or stored at 4 °C in the dark until image acquisition.

Confocal imaging of fixed samples was performed using an upright confocal microscope + Airyscan (LSM800, Zeiss) equipped with 2 GaAsP Photomultiplier Modules (PMTs) detectors using 40x/1.3 oil DIC, UV-IR objective. Multi-positions of Z-stacks (0.4μm step size) of fixed migrating cells were acquired using Zeiss software (ZEN 3.8).

### Primary and Secondary Antibodies

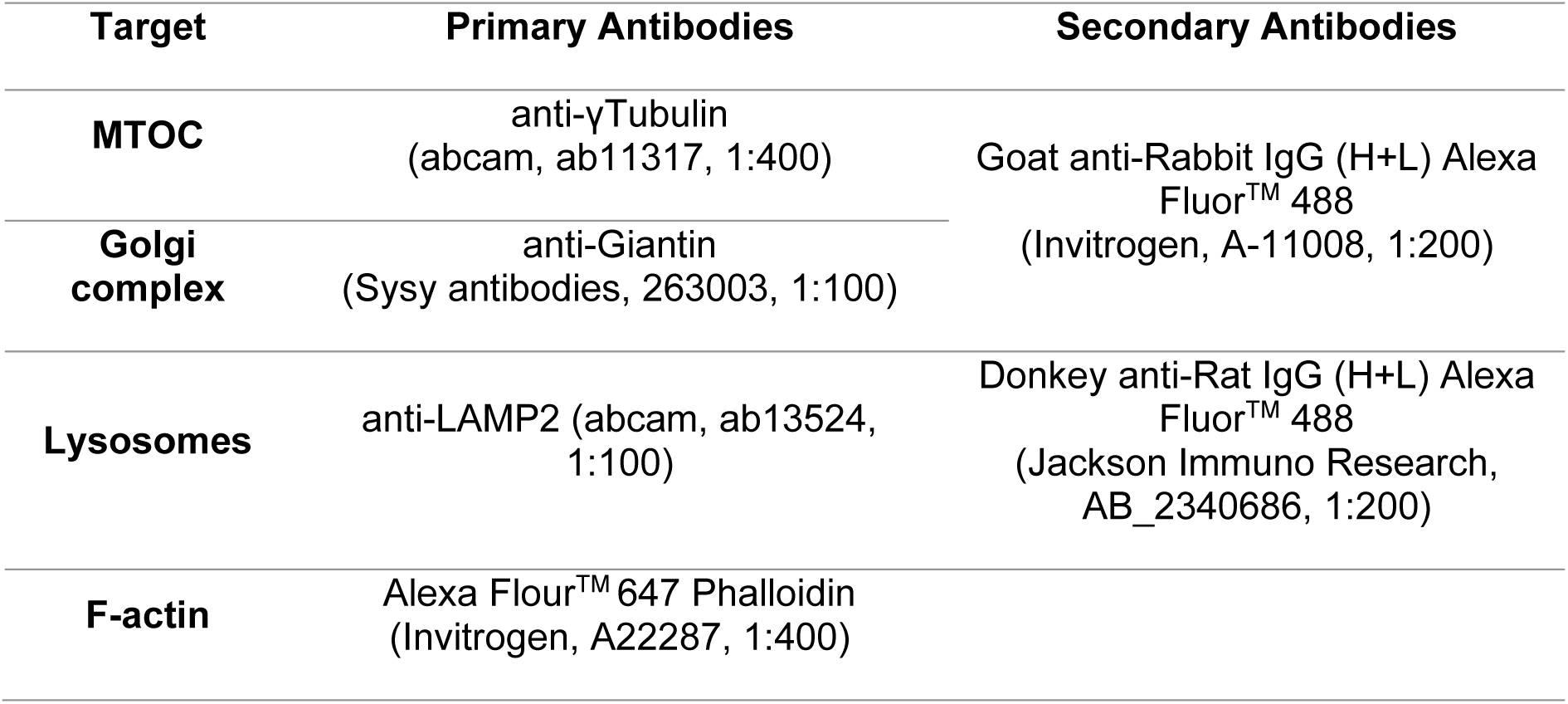

| Target | Primary Antibodies | Secondary Antibodies |
| --- | --- | --- |
| **MTOC** | anti-γTubulin<br>(abcam, ab11317, 1:400) | Goat anti-Rabbit IgG (H+L) Alexa Fluor™ 488<br>(Invitrogen, A-11008, 1:200) |
| **Golgi complex** | anti-Giantin<br>(Sysy antibodies, 263003, 1:100) |  |
| **Lysosomes** | anti-LAMP2 (abcam, ab13524, 1:100) | Donkey anti-Rat IgG (H+L) Alexa Fluor™ 488<br>(Jackson Immuno Research, AB_2340686, 1:200) |
| **F-actin** | Alexa Flour™ 647 Phalloidin<br>(Invitrogen, A22287, 1:400) |  |

### 2D analysis of cells migrating under agarose

For 2D analysis of cells migrating under agarose, LifeAct-eGFP expressing DCs labeled with Hoechst (NucBlue^TM^, Hoechst 33342, Invitrogen) were imaged with an inverted widefield Nikon TiE2 microscope equipped with 20x/0.5 NA PH1 air objective using a Hamamatsu EMCCD C9100 camera and a Lumencor Spectra X light source (390 nm, 475 nm, 542/575 nm; Lumencor). Images were taken every 30 seconds at multi-positions using the NIS Elements software (Nikon Instruments).

Single cells moving under agarose and their central actin pool were segmented based on the LifeAct-eGFP signal using Ilastik pixel classification^47^ and tracked using Fiji - Trackmate^48^. Resulting tracks were manually curated and only non-interacting, well-isolated cells with tracks longer than 10 frames (5 minutes) were further processed. Cell speed was calculated using the center of mass of the cell, based on the outline generated by the segmentation. Comparison of the temporal cross correlation of two parameters (central actin intensity, cell area, cell speed) and test for the statistical significance of the temporal offset we used cross-correlation analysis^49^ with a custom-written MATLAB script (MATLAB, R2020a).

### Bead displacement in agarose

To track the forces cells exert on the agarose, polystyrene microspheres with a nominal diameter of 1 μm and labeled with a fluorescent red dye (Red-580/605, F-13083 Invitrogen) were added to the agarose solution (1:100 dilution). The agarose cast and cell addition were performed as mentioned above.

Imaging of LifeAct-eGFP expressing cells labeled with Hoescht (NucBlue^TM^, Hoechst 33342, Invitrogen) was performed under physiological conditions using custom-built climate chambers (37 °C, 5% CO_2_, humidified) on an inverted spinning-disc confocal microscope (Nikon CSU-W1) cameras using a 40x/1.15 water objective. Z-stacks (0.1 μm step size) of migrating cells were acquired using two teledyne photometric BSI (USB3) sCMOS cameras with 95% quantum efficiency and a 6.5×6.5 μm pixel area. Images were acquired every 30 seconds for approximately 20 minutes.

Bead displacement analysis was performed using custom Python scripts. First, beads were individually segmented and labeled using their maximum intensity projection in Z and time to discard non-stationary beads, and a size filter was used to exclude bead aggregates. Next, we defined a fixed volume around each bead spanning the whole z-stack and 5×5 pixels in XY. In order to track the movement in Z, we generated a time kymograph of the bead intensity projected along the x and y-axis and tracked the moving bead edge, detected using Otsu threshold method **(Extended Data Fig. 2a)**. We then computed the total actin and nuclear intensity in each time frame within a similar volume of 20×20 pixels in XY, centered around each bead. In order to classify the actin contribution between none, cytoplasmic and central actin, we run a K-means clustering algorithm with two (DOCK8^−/−^ DCs) or three (WT DCs) clusters on the actin intensity curves. Hoechst (blue) signal was used to classify the nuclear contribution to the displacement. We then computed the average position of the bead for each of the regions and subtracted it from the baseline (no cell).

### Manufacturing and migration assays in microfabricated polydimethylsiloxane (PDMS)-based devices

#### Height Confiners

The microfabricated PMDS devices used to confined the cells in environments with different heights consist of two glass coverslips spaced by PDMS micropillars. One of the glass coverslips (#1.5, 22×22 mm, Mentzel, Thermo Fisher Scientific) was glued to a petri-dish with a 17 mm diameter hole using aquarium sealant, while the other, containing the PDMS micropillars, was attached to a PDMS cylinder which is secured by a magnetic device.

The pattern mold was produced by photolithography on a silicon wafer. The wafer was coated with SU8-GM1050 (Gersteltec) and soft-baked for 1 minute at 120 °C, followed by 5 minutes at 95 °C. The wafer was developed in SU8 developer for 17 seconds and then silanized with trichloro(1H,1H,2H,2H-perfluorooctyl)silane in a vacuum desiccator for 1 hour.

To produce the PDMS piston, silicone elastomer and curing agent were mixed in a 30:1 ratio, degassed as described before, and poured into an aluminium mold with the required dimensions. The PDMS pistons were cured at 80 °C for 6 hours and peeled off the silicon wafer with isopropanol.

Micropillars were produced by mixing silicone elastomer and curing reagent (PDMS Sylgard 184 Elastomere Kit, Dow Corning) in a 7:1 ratio. The mixture was then degassed using a planetary centrifugal mixer (ARE250, Thinky) and carefully poured onto the wafer. Round coverslips (#1.5, 10 mm diameter, Mentzel, Thermo Fisher Scientific) were plasma activated for 2 minutes (Plasma Cleaner, Harrick Plasma) and placed on the wafer with the activated surface facing the elastomere/curing agent mixture. The wafer was cured on a heating plate for 15 minutes at 95 °C, and the micropillar-coated coverslips were removed with a sharp razor blade and isopropanol.

The confiner devices were assembled by mounting a micropillars-bearing coverslip onto the PDMS piston with the micropillars facing upward and stuck to the glass bottom of the magnetic device. R10 medium was added to the micropillars-bearing coverslip and the petri dish containing the second glass coverslip, and incubated at 37 °C and 5% CO_2_ for at least 30 minutes to equilibrate. Matured DCs were resuspended in R10 with 2.5 μg/mL of CCL19 (PrepoTech) in a final volume of 20 μL and added to the micropillars. The PDMS piston with the micropillars and the cell mixture were then pressed onto the glass coverslip in the petri dish and sealed by a metal ring. Confined cells were incubated for at least 1 hour at 37 °C and 5% CO_2_ before imaging.

Imaging was performed as described in “bead displacement under agarose”. Z-stacks (0.4 μm step size) of migrating cells were every 30 seconds for approximately 20 minutes.

#### Pillar mazes, straight and constricted channels

The microfabricated PDMS devices containing pillar forest, straight, or constricted channels consist of PDMS blocks (fabricated as above, but using a 1:10 elastomer to curing agent ratio) attached to one glass coverslip. The devices were then cut in small squares (approximately 1×1 cm^2^) and attached to plasma-cleaned coverslips (#1.5, 22×22 mm, Thermo Fisher Scientific), and incubated at 85 °C for 1 hour.

The coverslips with the PDMS devices were then glued to a petri dish containing a 17mm diameter hole using aquarium sealant. Before adding the cells, devices were flushed and incubated with R10 medium for at least 1 hour at 37 °C and 5% CO_2_. 0.5-1×10^6^ matured DCs were added to one side of the devices and R10 with 2.5 μg/mL of CCL19 (PrepoTech) was added to the opposite side. Cells were incubated for at least 1 hour before image acquisition.

#### Analysis of the central actin pool intensity changes in cells moving in pillar mazes

Widefield images of LifeAct-eGFP expressing DCs moving in pillar mazes were acquired as described in “2D analysis of cells migrating under agarose”. Images were taken every 30 seconds at multi-positions with NIS Elements software (Nikon Instruments).

Cell area and nucleus were segmented and tracked employing the Ilastik pixel classification/cell tracking workflows. Non-interacting, well-isolated cells were identified and stabilized to their center of mass. For each cell, all regions of interest were manually annotated. The F-actin intensity in these regions was normalized to the overall F-actin intensity. All retraction events were pooled by shifting the events relative to each other such that t=0 marks the beginning of the retraction event and by setting the intensity in all regions to 1.

#### Analysis of cells moving in straight and constricted channels

Imaging of EB3-*mCherry* and LifeAct-eGFP expressing DCs in straight and constricted channels was performed in an inverted widefield Nikon TiE-2 microscope equipped with 40x/0.95NA DIC air objective using a Nikon DS-Qi2 CMOS monochrome camera and a Lumencor Spectra X light source (390 nm, 475 nm, 542/575 nm; Lumencor). Images were taken every 60 seconds at multi-positions with NIS Elements software (Nikon Instruments).

Actin distribution in cells during constriction passage was quantified using custom scripts in Python. First, channels were segmented using the bright field images, and the actin signal was averaged vertically (y-axis, only in segmented areas) in order to create a longitudinal actin density profile for each time frame. A maximum projection of these profiles resulted in a final time-averaged actin density profile, which was used to compute the ratio between the actin signal inside and outside the constriction.

#### Collagen migration assay of mature dendritic cells

Custom-made migration chambers were assembled using a petri-dish with a 17mm diameter hole in the middle which was covered by 2 glass coverslips^50^ (#1.5, 22×22 mm, ThermoFisher Scientific).

The collagen mixture, consisting of either 1.5 or 3mg/mL bovine collagen I (PureCol, Nutragen; both AdvancedBioMAtrix), was reconstituted by mixing 1.5-3.0×10^5^ matured DCs in suspension (R10 medium) with collagen I solution buffered to physiological pH with Minimum Essential Medium (Sigma) and sodium bicarbonate (Sigma) in a 1:2 ratio. In the experiments where labeled collagen was used, a mix of unlabeled and labeled collagen was used on a ratio of 1:2.

The collagen and cell mixture was then added to the migration chamber and allowed to polymerize in a vertical position for 1 hour at 37°C, 5% CO_2_. Directional cell migration was induced by overlaying the polymerized gels with 0.63μg/mL CCL19 in R10. To prevent drying out of the gels, chambers were sealed with paraffin (Paraplast X-tra, Sigma).

Brightfield movies were acquired in inverted cell culture microscopes (DM IL Led, Leica Microsystems) using either a 10x/NA or a 40x/NA air objective equipped with cameras (ECO415MVGE, SVS-Vistek) and custom-built climate chambers (37 °C, 5% CO_2_, humidified). Images were acquired with a time interval of either 30 or 60 seconds and global y-displacement was analyzed by a custom-made tracking tool.

#### Collagen fiber displacement and F-actin accumulation analysis

To visualize collagen fibers, collagen was directly conjugated to Alexa Fluor 594 NHS Ester (Succinimidyl Ester, ThermoFisher Scientific). Collagen was added to SnakeSkin Dialysis Tubes, 10K MWCO, 16 mm (ThermoFisher Scientific), and immersed in 100 mM NaHCO3 overnight at 4 °C to allow polymerization. Alexa Fluor 594 NHS Ester (1.5 mg/mL) was added to the polymerized collagen and incubated for 3 hours. To remove the unconjugated dye, the collagen mixture was placed in 0.2% acetic acid in deionized water for further dialysis overnight at 4 °C. Labeled collagen was kept at 4 °C until usage.

Movies of LifeAct-eGFP expressing DCs in Alexa-594-labeled collagen matrices were acquired on an inverted spinning-disc confocal microscope (Andor Dragonfly 505) using a 60x/1.4 NA objective and 488/561 nm laser lines in a custom-built climate chamber (37 °C under 5% CO_2_). Z-stacks (1.5 µm step size) of migrating cells were recorded using an Andor Zyla camera (4.2 Megapixel sCMOS) every 60 seconds for 20 to 25 minutes.

Collagen fiber displacement was calculated using the software Davis 8 (Lavision) applying Particle Image Velocimetry (PIV) as described before^20^. Computation of the closest distance between a collagen fiber deformation maxima and an F-actin intensity maxima across multiple time-lapse images and z-slices was performed using a standardized Python function. Briefly, this function independently finds the local maxima of collagen fiber deformation or F-actin intensity within a predefined neighborhood radius and calculates the minimum distances between these two structures per Z-slice and time point.

#### Fixation and immunofluorescence of collagen matrices

To visualize the nucleus-MTOC orientation during migration in different collagen matrices eGFP-Centrin expressing DCs labeled with Hoechst (NucBlue^TM^, Hoechst 33342, Invitrogen) were seeded in the collagen mixture and collagen gels were cast as described above. Three hours after the introduction of the CCL19 gradient, the collagen gels were isolated and immediately bathed in a PBS solution with 4% paraformaldehyde for 10 minutes at room temperature. The fixed collagen gels were washed with PBS at least three times and incubated with Phalloidin-Atto647N (1:400 dilution, Sigma) diluted in PBS supplemented with 0.2% BSA and 0.05% saponin for 2 hours at room temperature. After three more washes with PBS, the gels were mounted using Flourmount-G mounting medium with DAPI (00-4959-52, ThermoFisher Scientific).

Imaging was performed in an inverted confocal microscope (LSM800 inverted, Zeiss) equipped with 2 GaAsP PMTs detectors using a 40x/1.2 water objective. Multi-positions of Z-stacks (0.5 μm step size) of fixed cells migrating in the collagen matrices were acquired using Zeiss software (ZEN 3.8).

#### Statistics and reproducibility

Statistical details for each experiment can be found in the figure legends. Appropriate controls were performed for each biological replicate. All replicates were validated independently and pooled only when all showed the same trend. Statistical analysis was conducted in Prism10.2.2 (GraphPad). Data was tested for normal distribution using the D’Agostino Pearson Omnibus k2 test. Normally distributed data was tested using a student’s t-test or ANOVA. Non-normally distributed data was tested using the Mann-Whitney test. Categorical data (e.g. presence/absence of the central actin pool) was tested using Fisher’s exact test.

## Data availability

The authors declare that the data supporting the findings of this study are available in this manuscript. Data sets generated during this study are available from the corresponding author on request.

## Code availability

All custom-made scripts are available from the authors on request.

## Acknowledgements

This research was supported by the Scientific Service Units of ISTA through resources provided by the Imaging and Optics, Preclinical and Lab Support Facilities. In particular, we thank Mark A. Symth and Flavia G. G. Leite, from the Virus Service Team, who helped generating the lentiviral particles used in this study. We thank the Sixt group for valuable discussions and feedback. This work was supported by a European Research Council grant ERC-SyG 101071793 to M.S.. M.J.A. was supported by an HFSP Postdoctoral Fellowship LTF 177 2021 and A.J.G by a Lise Meitner Fellowship of the FWF (Austrian Science Fund). Y.F. was supported by the AMED-CREST (JP19gm1310005), the Medical Research Center Initiative for High Depth Omics, and CURE:JPMXP1323015486 for MIB, Kyushu University.

## Author contributions

P.R.-R., A.J.G, K.V, N.C. and M.S conceived the experiments. P.R.-R. performed and analyzed experiments with the help of N.C. and M.J.A.: N.C. performed and analyzed MTOC-nucleus position in microfabricated channels and under agarose; M.J.A. performed and analyzed bead displacement in agarose. F.G. provided data of DCs migrating under different agarose stiffness. M.R. wrote image analysis scripts for quantification of the collagen I fiber deformation and actin bursts proximity. J.M. generated microfabricated channels and pillar arrays. R.H. wrote image-analysis scripts for the analysis of the central actin pool in cells moving in pillar mazes, and cross-correlation analysis of central actin pool and cell area/cell speed. Y.F. provided reagents, technical support and advice. P.R.-R., M.J.A. and M.S. wrote the original draft and all authors critically reviewed the manuscript.

---

**Extended Data Figure 1 - related to Figure 1.**
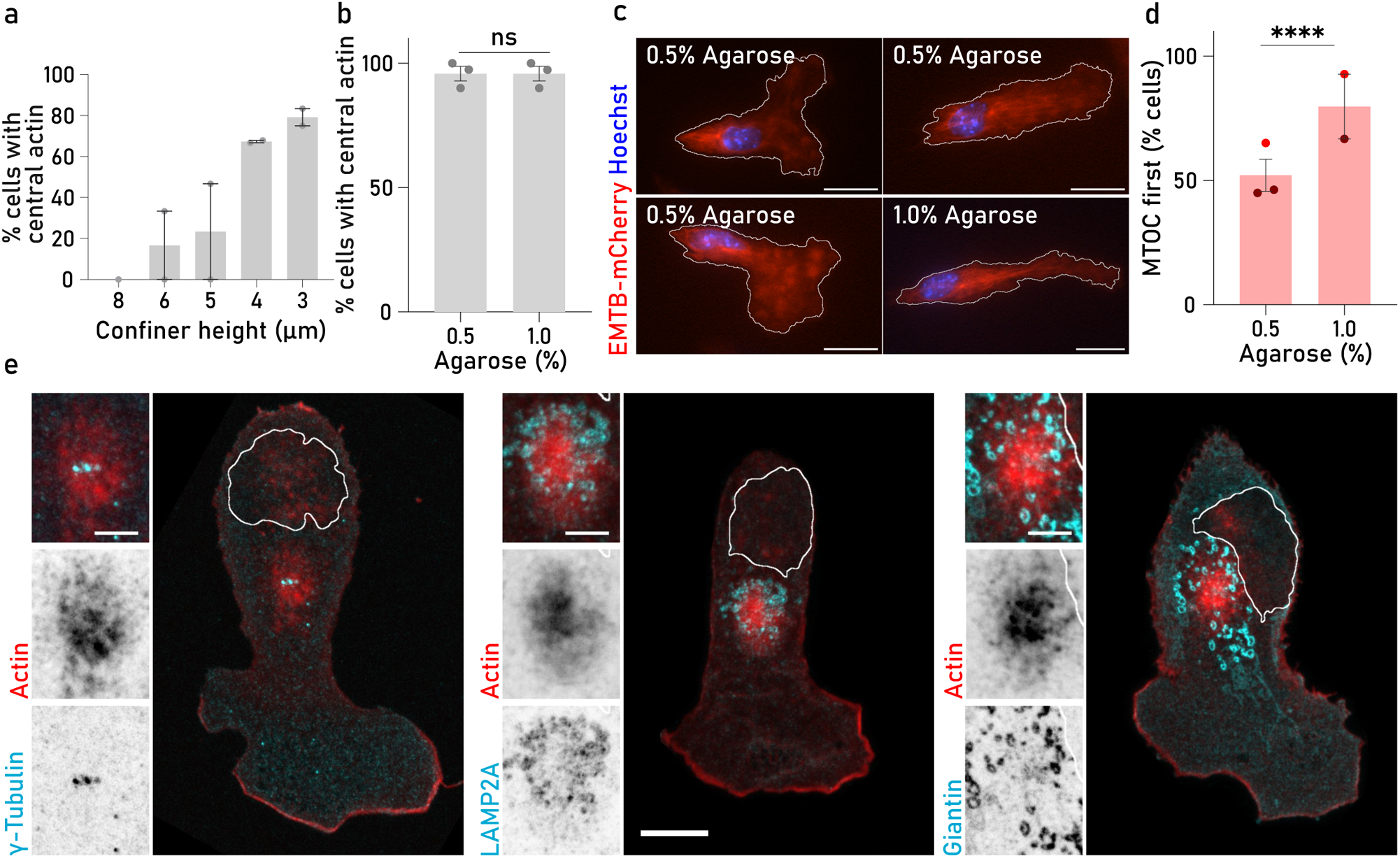
**a.** Cells showing the central actin pool under different height confinement (from 8 to 3 µm). Each dot corresponds to the percentage of cells per experiment where a central actin pool was detectable. 8 μm, n=10; 6 μm, n=63; 5 μm, n=17; 4 μm, n=67; 3 μm, n=14. Error bars show the SEM. **b.** DCs migrating under 0.5% or 1% agarose showing a detectable central actin pool. Each dot corresponds to the percentage of cells observed for three independent experiments. Cells migrating under 0.5% agarose, n=145; cells migrating under 1% agarose, n=88. Error bars show the SEM. ns p-value =0.4306. Fisher’s exact test. **c.** EMTB-*mCherry* (red) expressing DCS labeled with Hoechst (blue) migrating under 0.5% and 1% agarose. Note the different positionings of the MTOC in relation to the nucleus in 0.5% agarose. Scale bar, 20 μm. **d.** Cells showing MTOC-first orientation when migrating under 0.5% or 1% agarose. Each dot corresponds to the percentage of cells expressing either EMTB-*mCherry* (dark red) or eGFP-Centrin (red), where MTOC-first orientation was observed in at least two independent experiments. Cells migrating under 0.5% agarose, n=200; cells migrating under 1% agarose, n=74. Error bars show SEM. **** p-value<0.0001. Fisher’s exact test. **e.** Representative images of DCs migrating under 1% agarose and stained with Phalloidin (red) and antibodies specific for γ-Tubulin (MTOC - cyan, left), LAMP2 (Lysosomes – cyan, middle) or Giantin (Golgi complex - cyan, right). Nucleus position in these cells is shown by the white overlay. Scale bar, 10 µm. Scale bar inset, 3 µm.

**Extended Data Figure 2 - related to Figure 2.**
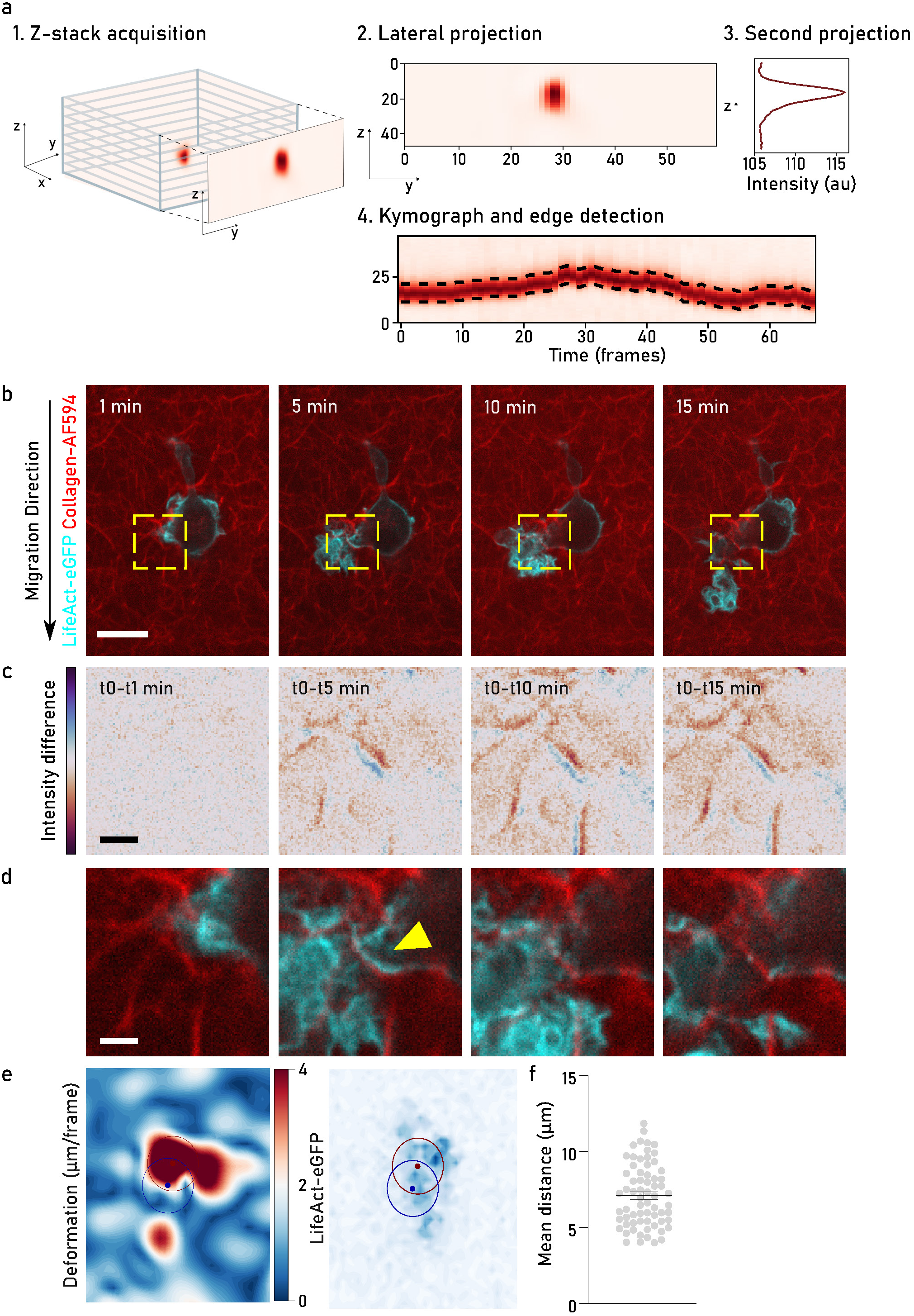
**a.** Workflow implemented for the quantification of bead displacement. **1.** Z-stacks acquisition of fluorescent beads embedded in agarose; **2.** lateral projection of the resultant Z-stack; **3.** intensity projection of the generated lateral projection; and **4.** kymograph and bead edge detection. **b.** Dendritic cells expressing LifeAct-eGFP (actin, cyan) migrating in collagen matrices labelled with AlexaFlour594 (red). Only one Z-plane is shown. The yellow dashed box shows the area for the insets in B and C. Scale bar, 10 μm. **c.** Collagen fibers in the area highlighted by the yellow box in B. Image subtraction of different time-points shows collagen fibers displacement over time. Red shows the fiber position in the initial time point while blue shows the position of the same fiber in the final time point. White represents areas where no changes in collagen fiber displacement were observed. Scale bar, 2 μm. **d.** Collagen fibers (red) and actin (LifeAct-eGFP, cyan) in the area highlighted by the yellow box in B. Yellow arrow shows actin accumulation at the place of collagen fiber deformation. Scale bar, 2 μm. **e.** Example of the workflow used for the quantification of the distance between collagen fiber displacement and actin accumulation. Maximum deformation of the collagen fibers and actin intensities in the different places of the Z-stacks acquired were calculated independently. Left: Collagen fiber deformation measured by Particle Image Velocimetry (PIV). Dark red shows high collagen fiber deformation while dark blue no deformation areas. Right: LifeAct-eGFP intensity. Dark Red dot shows the maximum deformation observed in the collagen fibers and dark blue dot the maximum intensity observed for LifeaAct-eGFP for this particular Z-slice. Distances in f were calculated between these maximum spots. **f.** Mean distances observed between maximum collagen fiber deformation spots and maximum actin accumulation spots. Each dot corresponds to the mean distance observed in each z-slice. n=10, error bars show SEM.

**Extended Data Figure 3 - related to Figure 3.**
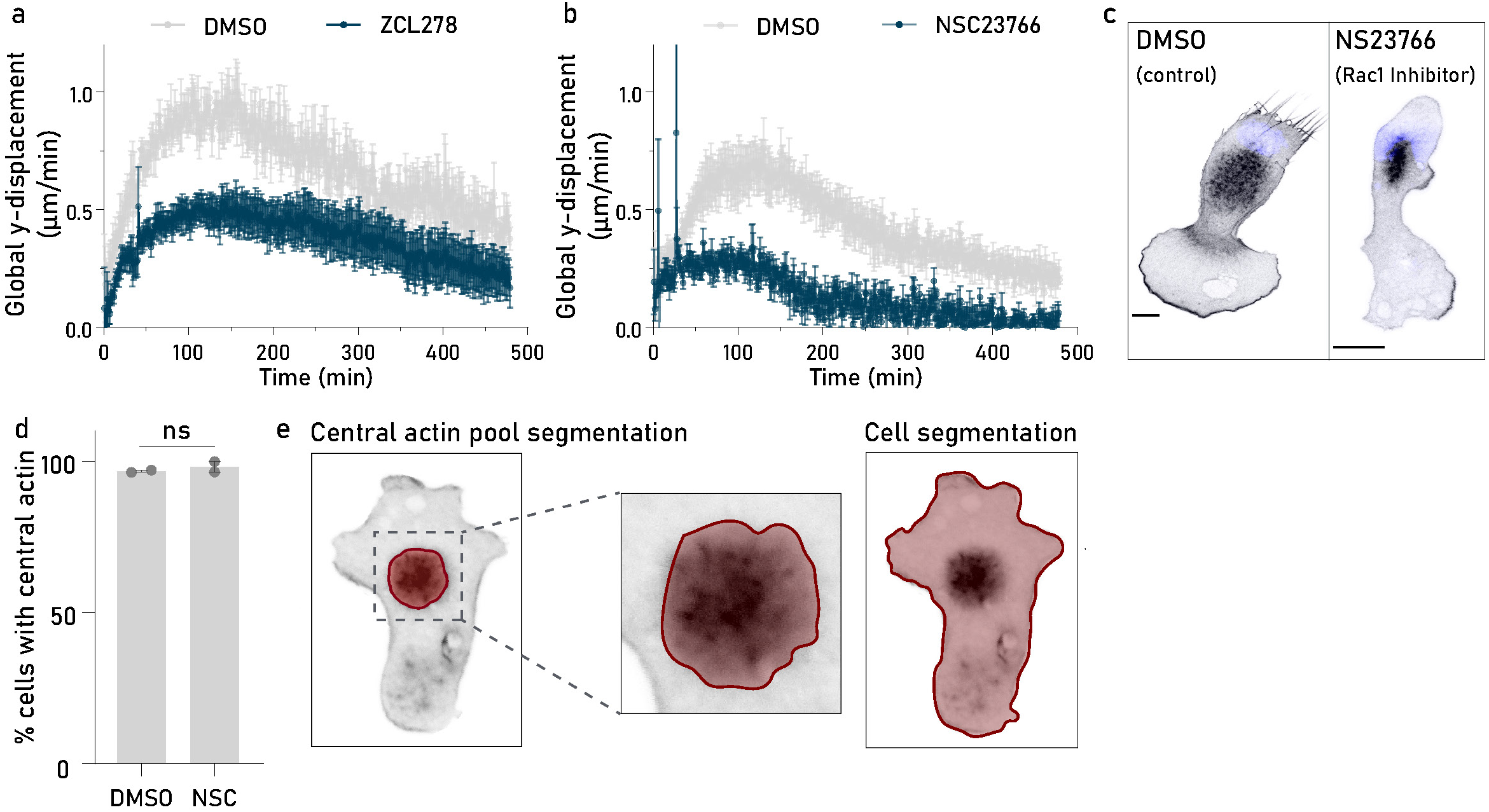
**a.** Global y-displacement of control (DMSO - gray) and ZCL278 (Cdc42 inhibitor, blue) treated DCs migrating in 1.7 mg/mL collagen gels. Each dot corresponds to the mean global speed in direction to the chemokine source observed in, at least, six different movies from three independent experiments. Error bars show the SEM. **b.** Global y-displacement of control (DMSO - grey) and NSC23766 (Rac1 inhibitor, blue) treated DCs migrating in 1.7 mg/mL collagen gels. Each dot corresponds to the mean global speed in direction to the chemokine source observed in, at least, six different movies from three independent experiments. Error bars show the SEM. **c.** DCs migrating under a patch of 1% agarose treated either with DMSO (control - left) or NSC23766 (NCS, Rac1 inhibitor - right). Cells were fixed and stained with phalloidin (F-actin, black) and DAPI (nucleus, blue). Scale bar, 10 μm. **d.** Cells showing a central actin pool upon treatment with DMSO or NSC23766 (NCS, Rac1 inhibitor). Each dot corresponds to the percentage of cells where a central actin pool was detectable in two independent experiments. DMSO, n=90; NSC (Rac1 inhibitor), n= 68. ns p-value>0.9999. Fisher’s exact test. **e.** Central actin pool was segmented based on its phalloidin intensity (left). Mean values obtained in the central actin pool region were normalized to overall F-actin intensity in the cell (right).

**Supplementary Figure 4 - related to Figure 4.**
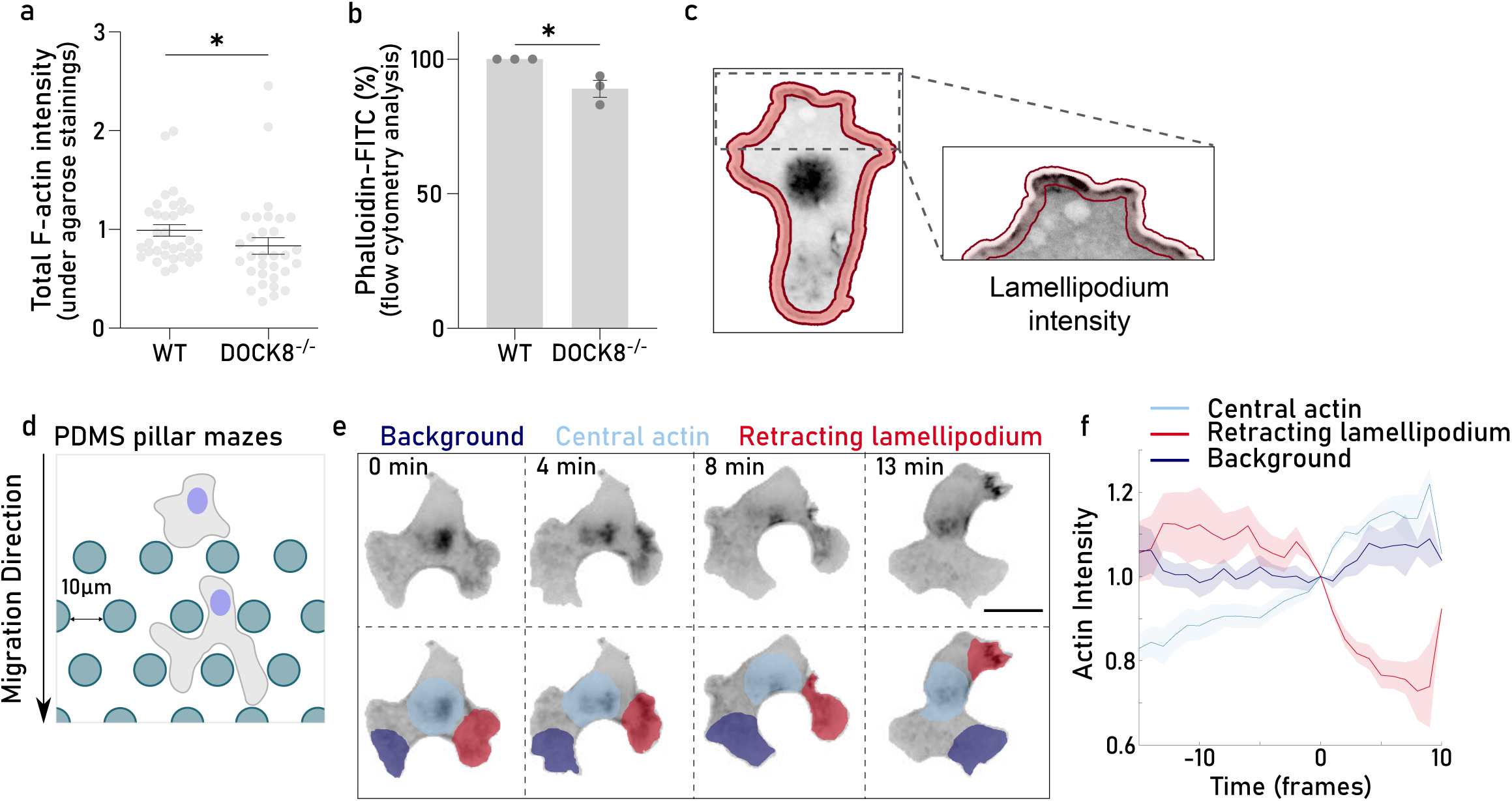
**a.** Total F-actin intensity (phalloidin) of WT and DOCK8^−/−^ fixed cells migrating under agarose. Each dot corresponds to the mean total F-actin intensity observed in one individual cell. Data was pooled from three independent experiments. **b.** Flow cytometry analysis of the total F-actin intensity of WT and DOCK8^−/−^ DCs fixed and stained with phalloidin-FITC. Each dot corresponds to the normalized F-actin intensity observed in the two cell types in three independent experiments. * p-value=0.03808. Paired t-test. **c.** Cells were segmented and the resultant outline dilated as shown by the red contour. For quantification of lamellipodial actin only the outline corresponding to the leading edge was considered, as showed by the inset. **d.** Scheme of cells migrating in a pillar maze. Cells were confined between surfaces 6µm apart and intersected by 10 µm distanced pillars. **e.** Top: Time-lapse images of a dendritic cell expressing LifeAct-eGFP (actin, black) migrating in a pillar maze. Scale bar, 15 µm. Bottom: Highlights of the three different regions of interest used for the analysis - retracting lamellipodium (red), central actin pool (light blue), and an area where no significant change in cell morphology was observed - background (dark blue). Scale bar, 10 μm. **f.** Actin intensity in the central pool (light blue), the retracting lamellipodium (red), and the background (dark blue) through time. Actin (LifeAct-eGFP) intensity in the regions of interest was normalized to 1 for time-point 0. Time point 0 was manually curated and corresponds to the time where lamellipodium retraction starts. n=24. Shadow error bars show standard deviation.

## Supplementary Videos Legends

**Supplementary Video 1:** Confinement induces amoeboid to mesenchymal transition and polymerization of a central actin pool in migrating dendritic cells (DCs).

*First part:* Epifluorescence movies of EB3-*mCherry* (red, MTOC), LifeAct-eGFP (black, actin) expressing DCs, labeled with Hoechst (blue, nucleus) migrating in PDMS channels with a narrow constriction (1.7 µm) at the entrance. *Second part:* LifeAct-eGFP (black, actin) expressing DCs migrating in confiners of different heights: 8 µm (top) and 4 µm (bottom). *Third part:* EMTB-*mCherry* expressing DCs (red, MTOC) migrating under soft (0.5%) or stiff (1%) agarose. Scale bar, 20 µm. *Fourth part:* LifeAct-eGFP cells (black, actin) migrating under soft (0.5%) or stiff (1%) agarose.

**Supplementary Video 2:** The central actin pool induces substrate deformations.

*First part:* LifeAct-eGFP DC (black, actin) labeled with Hoechst (blue, nucleus) migrating under agarose with beads labelled with AF-555 (red). *Second part:* Lateral projection of the cell showed before.

**Supplementary Video 3:** Local collagen fiber displacement is associated with actin bursts.

*First part:* LifeAct-eGFP expressing DC (cyan, actin) migrating in a collagen matrix labeled with AF-594 (red). White square shows the inset used for the close-up in the second part. *Second part:* close-up of the collagen fiber deformation (left) and the actin localization (right).

**Supplementary Video 4:** Low concentrations of Cdc42 and Rac1 inhibitors induces a slight decrease of cell speed in collagen matrices.

Bright field movies of WT DCs treated with DMSO (control – left), ZCL278 (Cdc42 inhibitor – middle) or NSC23766 (Rac1 inhibitor – right) moving in a collagen I matrix.

**Supplementary Video 5:** Central actin pool communicates with leading edge actin

*First part:* LifeAct-eGFP expressing DC migrating in a PDMS pillar maze. While left panel shows the cell migrating, the right panel highlights the regions used for the quantification shown in Extended Data Fig.4 f. Red shows the retracting lamellipodium, light blue the central actin pool, and dark blue the background. *Second part:* WT (top) and DOCK8^−/−^ (bottom) lifeAct-eGFP expressing DCs labeled with Hoechst (blue, nucleus) migrating under stiff agarose. *Third part:* LifeAct-eGFP expressing DC (top) and resulting segmentation of the cell body depicted in black, from which we extracted the cell area, and the central actin pool shown in red (bottom).

**Supplementary Video 6:** DOCK8 differentially affects DC locomotion depending on environmental factors

*First part:* Low magnification (10x) bright field movies of WT (left) and DOCK8^−/−^ (right) migrating in a collagen I matrix. *Second part:* High magnification (40x) bright field movies of WT (left) and DOCK8^−/−^ (middle and right) migrating in a collagen matrix. Middle panel shows and example of a DC fragmenting due to entanglement in the matrix. Right panel shows an example of a dying cell. *Third part*: WT (left) and DOCK8^−/−^ (middle and right) lifeAct-eGFP expressing DCs (black, actin) and labelled with Hoechst (blue, nucleus) migrating in complex PDMS pillar mazes. Middle panel shows an example of DOCK8^−/−^ cell with a single lamellipodium, while the right panel shows an example of a DOCK8^−/−^ cell forming multiple simultaneous lamellipodia during migration. *Fourth part*: Bright field images of WT and DOCK8-/- DCs labeled with Hoechst (blue, nucleus) migrating in straight PDMS channels.

